# SupeRJump: Determining normal and leukemic differentiation fate through semi-supervised jump diffusion modeling

**DOI:** 10.64898/2026.07.01.735284

**Authors:** Michael Bowman, Roopsha Bandopadhyay, Varsha Singh, Maria Telpoukhovskaia, Robert Vander Velde, Sydney M. Shaffer, Jennifer J. Trowbridge, Robert L. Bowman

**Affiliations:** Department of Cancer Biology, Perelman Cancer Center, University of Pennsylvania, Philadelphia, PA, USA; The Jackson Laboratory, Bar Harbor; ME, 04609, USA; Department of Pathology and Laboratory Medicine, Perelman School of Medicine, University of Pennsylvania, Philadelphia, PA, USA; Department of Bioengineering, School of Engineering and Applied Science, University of Pennsylvania, Philadelphia, PA, USA

## Abstract

Single cell RNA-seq (scRNA) has provided unprecedented resolution into cellular and clonal heterogeneity. Computational approaches have enabled recovery of differentiation dynamics, yet current approaches do not evaluate discontinuous differentiation processes present in malignant leukemia. To address these gaps, we developed SupeRJump: a jump-drift-diffusion based supervised cell-fate model (https://github.com/namwob44/SupeRJump/). We deploy this approach in human bone marrow, murine aging hematopoiesis, and lentivirally barcoded mouse models of acute myeloid leukemia. Our framework introduces a semi-supervised pseudotime strategy to fit a jump-drift-diffusion model and batch correction for lineage fate predictions from absorbing Markov chains. We introduce metrics to quantify a cell’s skewness toward particular lineages, transitions through intermediate progenitor states toward terminally differentiated states, and discontinuous transition dynamics. We use these metrics to identify cells preferentially biased for differentiation, their underlying transcriptional networks, and gene programs responsible for differentiation discontinuity.

## INTRODUCTION

Hematopoiesis is arranged in a hierarchy of stem and progenitor cells capable of producing distinct cell fates. Alterations in these differentiation processes is a critical feature underlying hematopoietic disease including myeloid leukemia. Understanding the cellular and transcriptional context for differentiation biases may offer insights into maximizing therapies that normalize hematopoietic function. Single cell RNA-seq (scRNA) has provided a critical function in exploring insights into these cellular processes.

Computational approaches have made recovering differentiation dynamics tractably possible. Such tools have been built on diffusion map paradigms introduced by destiny[1], and other diffusion strategies have adapted stochastic network traversal methods to reconstruct pseudotime[2] and infer high density transitory states[3]. New strategies have emerged to infer cellular outcome from earlier progenitor cells based on reformulating Markov chains with absorbing states[4],[5],[6],[7]. While scRNA does not uniformly sample all transition states in heterogeneous tissues, recent approaches have suggested highly sparse cellular states may represent unstable biological transition points[3]. Current approaches do not consider discontinuous differentiation processes often present in aberrant hematopoiesis such as leukemia. We aim to explicitly model discontinuous processes within the cell state transitions to account for traversal of under sampled sparse cell states.

To address these gaps, we developed SupeRJump: a jump-drift-diffusion based supervised cell-fate model. We present a hypothesis generation tool that identifies cell populations poised for differentiation and transcriptional networks underlying these transitions. We also introduce metrics to account for traveling to intermediate states on their way to terminal states. To demonstrate the effectiveness of our strategies, we deploy this approach on multiple datasets including: 1) human bone marrow [4], 2) normal murine, aged hematopoiesis [8], and 3) barcoded murine acute myeloid leukemia ex vivo cultures. We use these datasets to showcase different aspects of the approach. For each of these datasets, we identify cells with preferential bias for differentiation lineages, and their underlying transcription factor activity.

## RESULTS

### Model overview: SupeRJump enables robust lineage predictions

Our workflow is demonstrated in **Figure 1a**. After standard processing of scRNA we obtain labeled cell clusters. We provide a semi-supervised pseudotime that represents the expected ordering of cells. Afterwards, we fit our jump-diffusion model to build a cell-to-cell network. Our approach then aims to formulate the inference problem into two distinct phases, 1) inferring the differentiation outcome and 2) identifying transcription factor activities associated with differentiation. Regarding differentiation inference, we improve upon existing methods of cell-fate by accounting for discontinuous transitions in our cell-cell network using jump-diffusion, introducing batch correction to handle imbalances of heterogenous populations across replicates and conditions, and applying a semi-supervised signature as our pseudotime measure. With these improvements, we introduce a novel strategy that identifies and characterizes candidate cells that are preferentially biased toward particular lineages. In addition to calculating differentiation likelihoods for each cell to each lineage (fate probabilities), we also introduce metrics to account for intermediate state transitions termed ‘visitation probability’ and ‘weighted destination time’. To uncover potential mechanisms underlying lineage-specific differentiation, we deploy two primary strategies 1) analyzing gene ontology terms associated with discontinuous processes, and 2) multivariate linear modeling to evaluate lineage fates as a function of inferred transcription factor activity. Collectively we use these approaches as a hypothesis generation tool to nominate mechanistic drivers of a lineage bias and output.

**Figure 1:**
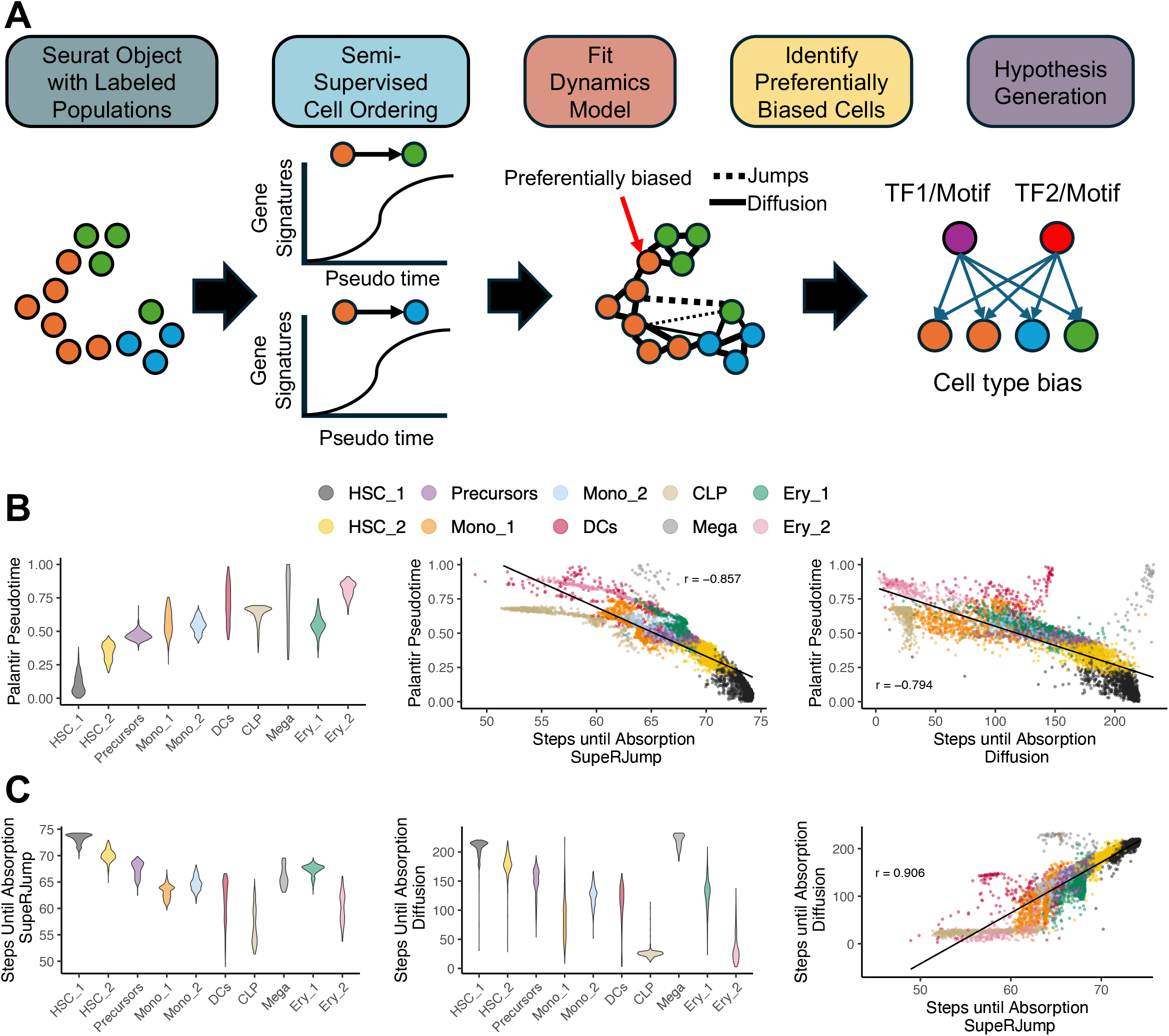
SupeRJump enables faster state absorption than strict diffusion A) Overall framework of SupeRJump approach. B) Violin plot depicting Palantir pseudotime (y-axis) and cell types (x-axis) from a human hematopoiesis dataset (Left). Scatterplots depicting the correlation between Palantir pseudotime (y-axis) and steps to absorption for strict diffusion model (middle; r = −0.794) and jump-diffusion model (right; r = −0.857). C) Violin plots depicting steps to absorption (y-axis) for SuperRJump (left) and strict-diffusion (middle) across cell types (x-axis). Diffusion only strategies have heavier tails suggesting they have restricted neighbors and do not capture discontinuous processes that occur. Scatterplot depicting correlation between steps to absorption for SupeRJump (x-axis) and strict diffusion (y-axis; r=0.906)

### Jump incorporation accurately reflects pseudotime ordering

Current diffusion policies implement k-nearest neighbors which limits the number of neighbors accessible to any given cell[7]. Our strategy expands the allowable neighbors through a probabilistic approach via jump-diffusion model fitting. Expanding the neighbors for each cell broadens the range of possible transitions, allowing us to capture rare events. Consequently, this expansion allows for faster traversal of the cell-cell network. To directly compare transition probability matrices (TPMs) with and without the jump model, we analyzed a previously described human bone marrow dataset[4]. Both TPMs utilize a Palantir pseudotime[4] (**Figure 1b**) and the same absorbing cells from CellRank2[7] for fair comparison. These absorbing states allow us to calculate both steps to absorption, and fate probability. We compare performance by correlating the pseudotime with the number of steps it takes for a cell to reach an absorbing cell. The TPM with jumps has a stronger correlation (r = −0.857) between the absorption and pseudotime than the diffusion only (r= −0.794; **Figure 1b**).

We next assessed the distribution of absorption steps by cell type (**Figure 1c**). We observed that compared to the jump TPM, the diffusion TPM had heavier tailed distributions which can be a consequence of limited transition options in a constrained diffusion only model with fixed nearest neighbors (kurtosis HSC_1 for diffusion only model: 26.44, jump-diffusion: 0.6). This can result in few cells serving as transitory states towards terminally differentiated states, and a relative increase in steps to absorption for cells more disconnected from highly connected transition states. Jump diffusion models allow for a dynamic increase in neighbors for each cell, mitigating the over-reliance on rare, high transitory cells. Our approach correctly orders the erythroid and myeloid lineages simultaneously, rather than sequentially, with monocytes, megakaryocytes and early erythroid cells being absorbed at a similar number of steps. Finally, we demonstrate the two strategies are strongly correlated (r=0.906), which suggests both models are primarily following similar dynamics.

In addition to quantifying steps to absorption, we next sought to understand which clusters are more likely to be visited on their way to a terminally differentiated state, a metric we termed ‘visitation probability’. In brief, we establish an absorbing Markov Chain and use the fundamental matrix to derive a metric from a first-passage analysis (as outlined in Methods). We then use a heatmap to visualize the aggregated visitation probability (color scale) of cell types (rows) transitioning to other cell types (columns) before absorption (**Figure 2a**). We observed that jump models allow greater branching while diffusion strategies are more isolated within a single cell type. Nuances between these two models highlight how network connectivity plays a crucial role in determining which progenitor path cells are likely to take for lineages. In the diffusion approach, we identified that HSC-1 is a more likely branch point between myeloid and megakaryocyte (MK) lineages (HSC-1àMK = −0.07; HSC-2àMK= −0.49). Yet in the jump model, HSC-2 carries greater MK visitation probability than HSC-1 (HSC-1àMK = −0.18; HSC-2àMK = 0.35). We perform an assortativity measure (bounded between −1 and 1) on each network to identify intra-cluster (1) vs inter-cluster (−1) connections and find the diffusion model is 0.75 while the jump model is 0.33. By incorporating jump-models, we allow for more branching, capturing unobserved phenomena in the diffusion setting.

**Figure 2:**
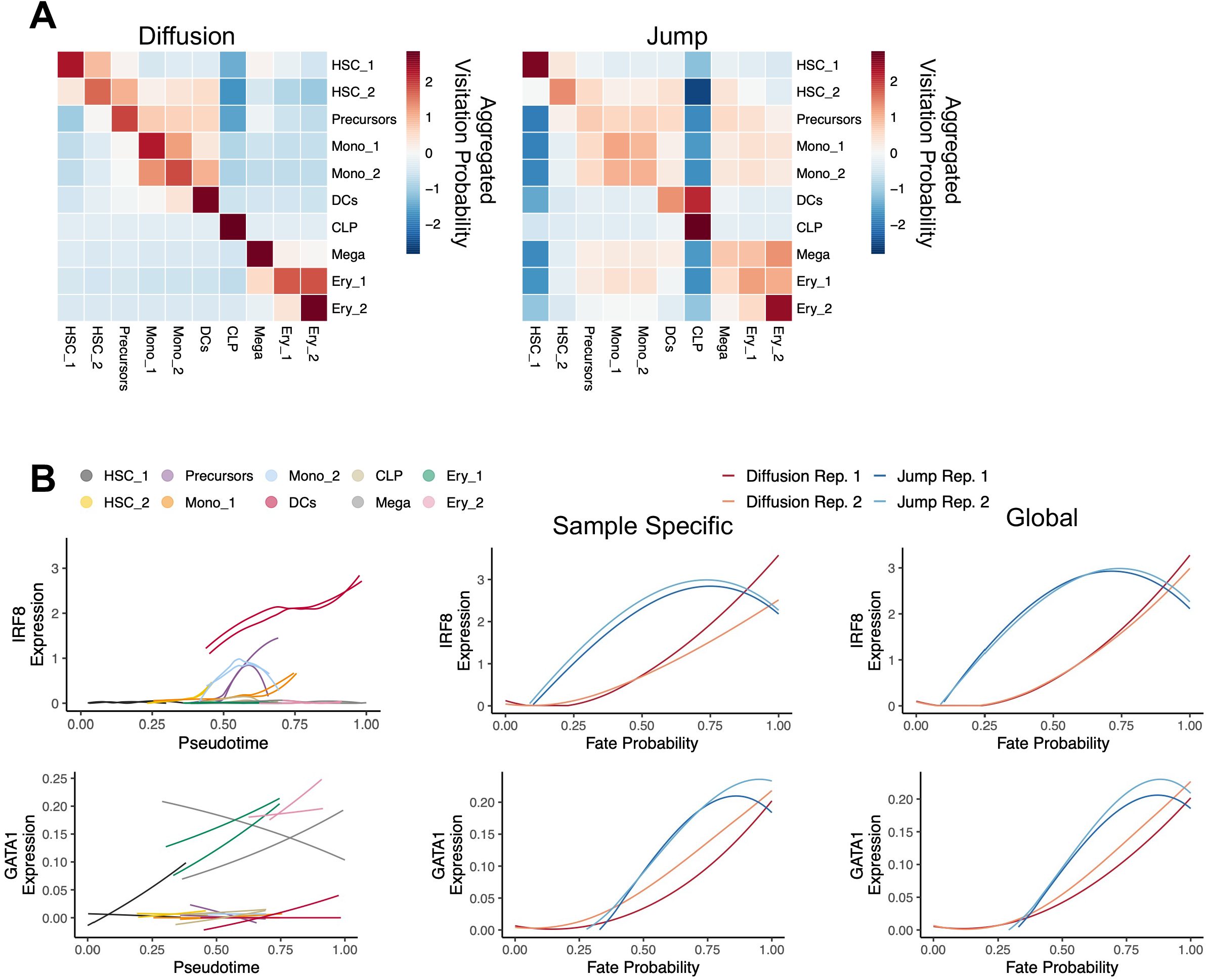
SupeRJump network differences revealed through visitation probability and fates A) heatmap for aggregated visitation probability for the Diffusion only, and SupeRJump models. Rows are initiating state and columns are destination state. B) line plots depicting either pseudotime or fate probabilities (x-axis) by RNA expression (y-axis). Multiple lines represent replicates. The top row is for *IRF8* expression and the bottom row shows the *GATA1*. The middle panels are for individual TPMs while the right column is the pooled TPMs.

In most approaches, TPMs are constructed by pooling all replicates and conditions into a single network, ultimately limiting the statistical power of unique biological replicates and failing to capture the heterogeneity of intermediate cell states across conditions. To account for cell composition heterogeneity, we employ a batch correction scheme inspired by a bulk-RNA seq strategy, ComBAT[9, 10]. This allows for sample specific fate probability determination, where we additionally restrict each biological condition to absorbing states represented in their own network, as opposed to across all samples. We returned to the data from Setty et al[4] to determine how batch corrected fate probabilities correlated with well-defined transcriptional programs. We evaluated lineage specific transcription factor expression by pseudotime for individual cell populations and samples (**Figure 2b**). Expectedly, we observed increased RNA expression levels of *IRF8* and *GATA1* for dendritic and erythroid lineages respectively. We next correlated expression levels with fate probabilities across all cell types and observed. IRF8 increases when probability for dendritic lineage increases, likewise the same trend occurs with *GATA1* along the erythroid lineage. In this setting, replicates displayed consistent behavior in fate probabilities for both the sample-specific and global model. We posit that this previously inaccessible comparison between replicates will be critical for comparison of fate probabilities in experimental settings with severely imbalanced cell populations or in less well-defined cellular systems such as leukemia, both of which are explored in **Figure 5**.

### Gene expression programs identify biological processes linked with jumps and differentiation fate

One unique benefit of jump-drift-diffusion modeling is the potential to identify underlying transcriptional programs associated with model fits that are best represented by discontinuous processes. Our strategy, which we call JumpROPE (Jump Relevant Ontology Pathway Extraction), enables hypothesis generation for discontinuous processes associated with transitions toward specific cell types. We fit a jump-model for each principle component-cell type pair, with a mean reversion to abstracted ‘eigen-cells’ (see Methods). The magnitude of the jumps corresponds to how large a deviation is needed within the principle component space to reach that cell type specific eigen cell (see Methods). To explore this approach, we fit the jump model on a scRNA dataset containing bone marrow cells from young and middle-aged mice from 14 biological samples (n=52967 cells)[8]. We then identified principle components with jumps toward specific cell types (Figure 3a). As an example, the 4^th^ PC depicts a jump where we identified discontinuous processes for cells toward the GMP population (Figure 3b). The increase in probability density after the initial dip demonstrates a jump occurs within the observed pseudotime 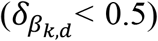. In contrast, we found a strictly diffusion model for the 29^th^ PC toward the GMP population, with a jump rate outside the observed pseudotime 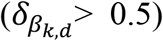. After identifying where jumps and discontinuities potentially exist, we extract genes which contribute most to this specific PC, split them based on their positive and negative loadings along the PC, and perform gene ontology on both subsets. JumpROPE results for PC4:GMP identified an association with cell cycle progression in the negative PC-loadings and transcription regulation in endoplasmic reticulum (ER_ stress in the positive PC-loadings, suggesting discordant regulation of these processes in GMP fate (Figure 3c). ER stress and secretory capacity in upstream multi-potent progenitor (MPP) cells has been linked to myeloid-erythroid skewing[11], with high secretory cells showing higher stress and lower long term engraftment capacity[12]. Overall, we identified that, instead of a common gene set, each model fit associated with a jump was enriched for relatively unique gene expression programs (Supplementary Table 1). These results suggest that the mechanisms underlying discontinuous differentiation processes might vary by cell type.

**Figure 3:**
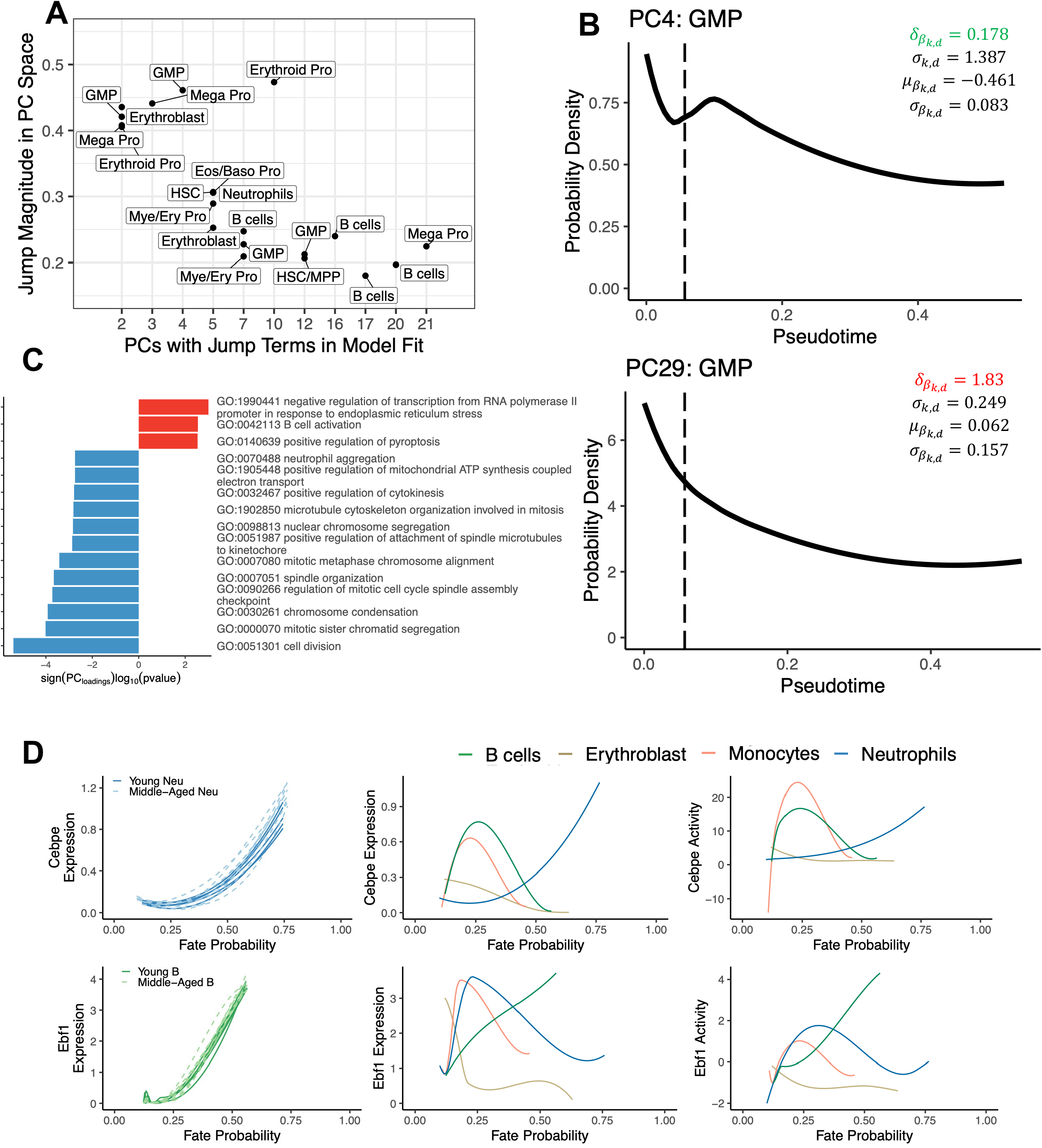
Linking biological processes to jumps and differentiation fate A) A scatterplot showing the jump magnitude identified for eigen cells (y-axis) along different principle components (x-axis). B) two probability density functions (PDF) on y-axis, along pseudotime (x-axis) for GMPs along PC4 (top) and PC29 (bottom). The dashed line indicates the eigen cell pseudotime. A jump occurs, if the model fit chooses a 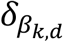 within the maximum pseudotime observed. In this instance, 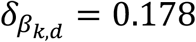 which is less than our maximum pseudotime. The bottom plot shows a characteristic of diffusion model fits with a smooth decaying since the 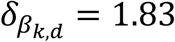 is beyond the maximum pseudotime. C) A barplot for the topGO terms associated with PC4. Negative loadings are blue, while positive loadings are in red. D) lineplots comparing batch corrected fate probability (x-axis) and RNA expression (y-axis) for each replicate within the dataset. The dashed lines indicate Middle-Aged while the solid lines indicate Young replicates. The top row is for *Cebpe* RNA and TF activity while the bottom row is for *EBF1*. Lineplots are also show in the middle and right columns with the fate probabilities (x-axis) across four lineages with their respective RNA or TF activity on the y-axis.

We next sought to evaluate how fate probabilities correlated with transcription factor (TF) networks. We batch corrected fate lineages to correct for replicates across conditions and compared the expression of lineage-specific transcription factors *Cebpe* and *Ebf1* across neutrophil and B cell lineages (**Figure 3d**). We observed a significant lineage-specific association between Cebpe and fates (F-value: 10422.7, p-value < 2.2×10^−16^) with post-hoc comparisons of lineage-specific slopes demonstrating specificity for the neutrophil lineage (p<0.001). We next evaluated TF activity using a multivariate linear model[13] and observed Cebpe TF activity also showed an expected lineage-specific association with fates (F-value: 42816.29, p-value < 2.2×10^−16^) with a post-hoc comparison again identifying neutrophil lineage as significantly different from the other lineages (p<0.001). Performing the same mixed effects model on Ebf1 and fates reveals similar corresponding trends with statistically significant B cell lineage specificity (p<0.001) with post-hoc correction. The full breakdown across both genes and transcription factors is available in **Supplementary Table 2**. Collectively, JumpROPE and TF activity results demonstrate a roadmap by which biologically meaningful information can be extracted from jump-diffusion modeling and fate probability evaluation.

### Uncovering preferential lineage bias in aging normal hematopoiesis

Cell cluster identification in scRNA data can often result in an oversimplification of cell type identity where broad classifications can encompass several distinct cell-types possessing unique differentiation capacities. This is particularly problematic in leukemia where mature cell type-specific gene expression programs are often blended with self-renewal programs present in earlier progenitors. Reference mapping to cellular atlases has offered increased resolution into these cell type assignments, but remain dependent upon expert curation from bulk gene expression data. We sought to complement these approaches by devising a fate probability-based approach for identifying biased cells within a broad cell population. We focused on GMPs, a heterogenous progenitor population capable of producing several differentiated cells including monocytes and neutrophils.

For each state-fate cell pair, our bias detection relies on aggregating weighted negative log-probabilities from: 1) a specific fate lineage (neutrophils), 2) membership of the lineage specific cell type (neutrophils), and 3) membership of the self-identified cell state (GMPs). We use this aggregated outlier score to account for skewness of a given GMP cell towards a given lineage. We then apply a secondary robust quantile regression filter (bias score) to identify additional biased cells that would have been masked by extreme skewing. Applying the cutoff on the outlier score directly yields n=134 cells, while applying the cutoff on the bias score yields n=1031 cells (**Figure 4a**). We next applied this bias metric to the neutrophil and monocyte lineages separately allowing us to identify whether GMP cells were unbiased, biased for one lineage, or bipotent (**Figure 4b**).

**Figure 4:**
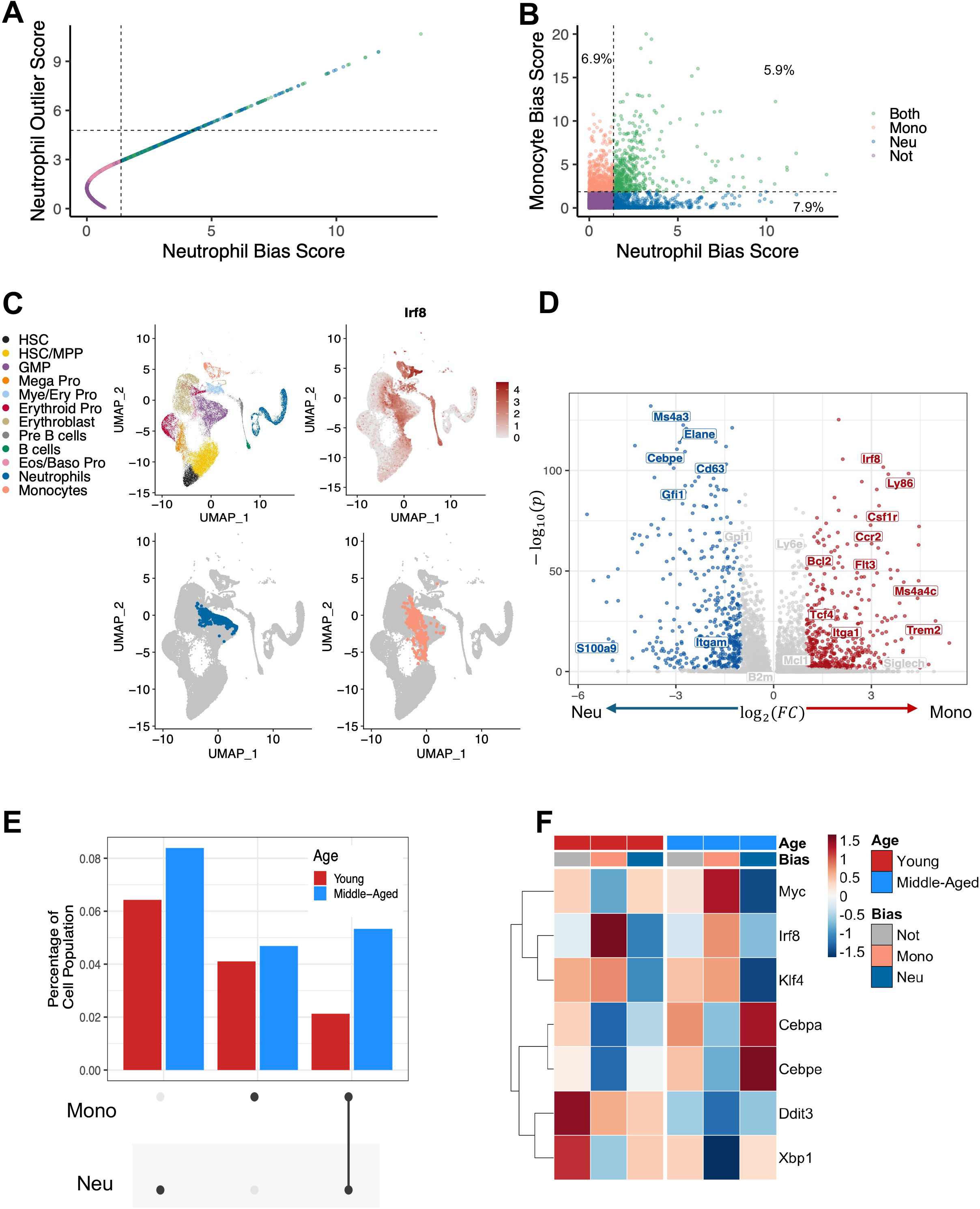
Preferential lineage bias provides cells of interest within populations A) A scatterplot for GMPs using our bias score for neutrophil lineages (x-axis) compared to a conventional outlier score (y-axis). The dashed lines represent the Tukey upper bound cutoff. B) A scatterplot comparing GMP cells for two lineages, Neutrophils (x-axis) and Monocytes (y-axis). The dashed lines indicate the Tukey cutoff. We achieve nonbiased, unibiased and multibiased GMP cells, with the percentage of cells falling into each bias category shown. C) shows a UMAP of the cell types, *Irf8* expression, and both uni-biased cells for their respective lineages. Monocyte biased GMPs appear to follow the column of *Irf8* expression. D) a volcano plot showing differential RNA expression on uni-biased GMPs for monocytes (red) and neutrophils (blue). E) an upset plot showing the proportion of GMPs cells biased for lineages split by their Age association (Young is red while Middle-Aged is blue). F) A heatmap of transcription factor activity for the unibias and nonbias GMPs split by age. Monocyte biased cells having increased TF activity for Irf8 and Klf4 while neutrophil biased have higher activity for Cebpa and Cebpe.

**Figure 5:**
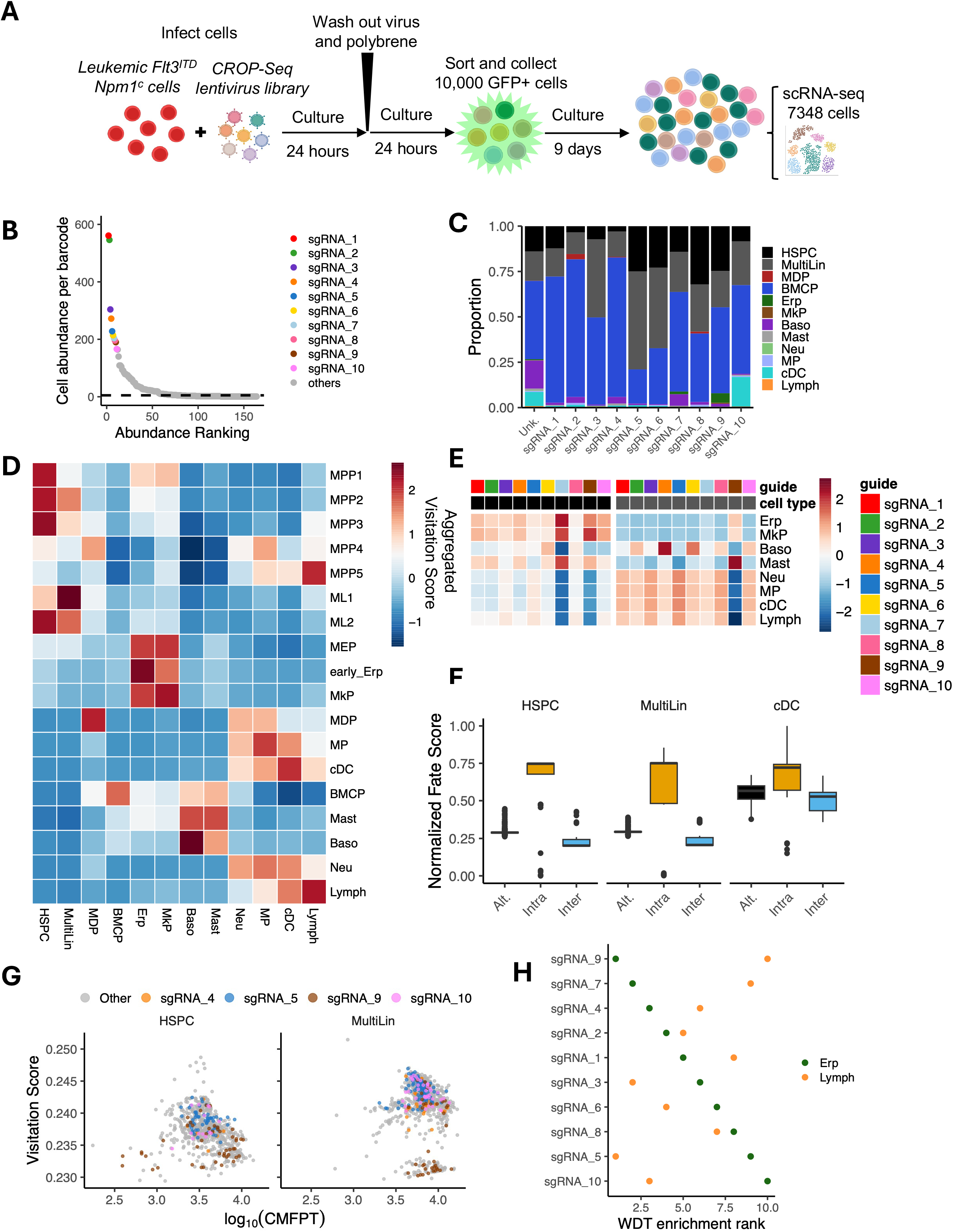
Lentiviral barcoding supports intraclonal fate transitions A) The experimental workflow of *ex-vivo* assay. B) a scatter plot for cell abundance of each identified guide ranked by the number of cells identified. The dashed line was a cutoff of 5 cells. The top 10 guides are colored. C) stacked barplot for cell type proportions across the top 10 guides identified and the aggregated population of unknown/unidentified sgRNA cells. D) A heatmap of the aggregated visitation scores for refined cell types (rows) navigating to broad clusters (columns). E) A fate heatmap broken up by HSPC and MultiLin cells that has moderately positive correlation with the proportion of cell types in C) (HSPC: *ρ*= 0.490, MultiLin: *ρ*= 0.495). F) boxplots for the normalized fate score for cDC lineage. Plots are split by cell type, and colored by connection type (black: Alternative, yellow: Intra-clonal, blue: inter-clonal). G) scatter plots split by cell types navigating toward the highest cDC fated MultiLin cell (a part of sgRNA_5) before reaching a terminal state. The x-axis is the time to reach the MultiLin, and the y-axis is the visitation score. Color shows the identified guide for easier visualization. H) a scatterplot comparing the Weighted Destination Time enrichment ranks of guides for two lineages (erythroid and lymphoid).

Differential expression analysis identified *Irf8*, a critical driver of monocyte and dendritic cell development, as one of the most enriched genes in monocyte-biased GMPs (n=515 cells) compared to neutrophil-biased GMPs (n=588 cells) (Figure 4cd). We additionally identified upregulation of Ccr2, Ly86, and Ly6e in monocytic-biased GMPs, confirming an enrichment of a monocytic-gene expression program in these cells. Conversely, neutrophil-biased GMPs possessed upregulation of expected genes including *Cebpe, Cd63, Elane, S100a9*, and *Gfi1* (**Figure 4d, Supplemental Table 3**). These findings were supported by geneset enrichment analysis of differentially enriched genes including monocytic (p<2.92×10^−56^) and dendritic (p<5.11×10^−60^) genesets enriched in monocyte-biased GMPs, meanwhile neutrophil-biased GMPs were enriched for genesets involved in neutrophilic degranulation (p<2.41×10^−15^; **Supplementary Table 4**).

Next, we assessed changes in lineage-biased GMPs as a result of aging, a process known to skew myeloid differentiation[13–17]. Specifically, we see neutrophil biased GMPs increase with aging (Young: n=130 6.4%, Middle-Aged: n=458 8.3%; p < 0.0039; **Figure 4e; Supplementary Table 5**). We next assessed age-specific transcriptional networks that changed amongst these biased-GMP populations. We observed expected enrichment for TFs that associate with specific lineages such as Klf4 and Irf8 for monocytic-biased GMPs while Cebpe and Cebpa were enriched in neutrophil-biased GMPs (**Figure 4f**). Lastly, we identified Ddit3 and Xbp1 had lower activity levels in middle-aged mice, nominating a role for dysregulated unfolded protein response in aging progenitor cells.

### Lentiviral barcoding supports intraclonal fate transitions

We next sought to experimentally validate fate probability predictions using lentiviral barcoding. We have previously developed a murine leukemia model driven by co-mutations in Npm1 and Flt3. Critically, this penetrant model of leukemia retains differentiation potential to a range of mature myeloid fates including monocytes and dendritic cells[18]. To assess clone-fate relationships, we infected whole bone marrow cells from leukemic mice with a CROP-seq barcoding lentivirus encoding GFP (**Figure 5a**)[19], [20]. After 48 hours, infected cells were sorted on GFP+ expression levels. Sorted cells were expanded for 9 days in liquid culture with pro-differentiation cytokine conditions (SCF+IL3) and then processed for single cell RNA sequencing (n=7348 cells). We identified 75 clones with >5 cells/clone (dashed line), with the top 10 clones representing 39.2% of cells in the dataset (Figure 5b). We recovered a range of cell types including primitive HSCs, MultiLin progenitors, classical dendritic cells (cDCs), baso-mast cell progenitors (BMCPs), and mature basophils, the latter two of which are common cell culture consequences of prolonged SCF+IL3 exposure (**Figure 5c; Supplemental Table 6**). We once again assessed visitation probability between clusters (**Figure 2a**), but this time evaluating a fine to broad cell type naming resolution to resolve intermediate states (**Figure 5d**). Despite the BMCP abundance, this sample and metric demonstrated canonical differentiation transitions with the MPP1/2 populations generally skewed for erythroid bias and MPP3s skewed for myeloid differentiation[21, 22]. The MPP4 population has potential to give rise to MDPs, MPs, and lymphoid lineage[21, 23]. MPP5 populations skew toward lymphoid lineages[24]. These examples highlight that visitation probability can recover progenitor transitions without the explicit over-interpretation of current fate predictions strategies.

This unbalanced cell type skewing amidst the high abundance of BMCPs presents a unique challenge to recover fate probabilities for relatively rare cell population such as cDCs. To investigate how clonal identity associated with fate probability, we focused on 4 clones: sgRNA-5 contains high proportions for HSPC (25%) and MultiLin (53.9%) populations, while sgRNA-4 contains the highest proportion of BMCP (76.8%). Both guides contain relatively rare cDC populations (sgRNA-4: 1.1%, sgRNA-5: 1.3%). The sgRNA-10 contains the highest proportions for cDC (16.2%). The sgRNA-9 contains no cDCs (0%), the highest erythroid progenitors (Erp: 5.2%), and a high HSPC (24%) population. We assessed lineage normalized fates in HSPCs and MultiLin cells for the 10 most abundant clones and found each lineage generally followed the distribution of cell compositions with significant positive correlation (HSPC: spearman *ρ* = 0.490, p< 1.2×10^−6^ MultiLin: spearman *ρ* = 0.495, p<9.12×10^−7^, **Figure 5e**). We observed that across all clones, as compared to MultiLin cells, HSPCs had a higher fate probability to the MkP and erthyroid lineages. This suggests the two separate progenitor populations are more likely to give rise to different lineages. We also observed clone specific behaviors with sgRNA-9 representing a higher MkP fate than other clones, mirroring the downstream cell type distribution for this clone.

Although fate correlated with cellular composition as expected, we next asked whether clones with similar cell-type distributions could nevertheless be distinguished by fate. Specifically, we sought to determine if cells have higher affinity for absorbing cells with their own clones than other clones. To demonstrate affinity within and between clones, we focused on the rare cDC compartment as a terminal absorbing state, which comprised cells from sgRNA-5, sgRNA-10, smaller subclones, and cells where an sgRNA could not be recovered (unknown). We considered three types of cell-cell connections based on transient and absorbing barcodes: 1) intra-clonal, when the transient and absorbing cells are from the same clone (e.g. HSPC from sgRNA-5 with a fate toward sgRNA-5 cDC), 2) inter-clonal, when the transient and absorbing cells have a mismatched clone (e.g. HSPC from sgRNA-10 with a fate toward sgRNA-5 cDC). and 3) alternative, similar to inter-clonal but the transient cell’s clone is not represented in the absorbing states pool (e.g. HSPC from sgRNA-4 with a fate toward sgRNA-5 cDC). To correctly classify connections, we filtered out all transient cells that had unidentified clones and then normalized cDC fate scores for three separate cell populations including HSPCs, MultiLins, and cDCs (Figure 5f). Across all three populations, intra-clonal connections have the highest fates, and inter-clonal connections were lowest across all three connection types, implying intra-barcode traversal is more likely to occur. (2-way ANOVA F-value=786.73, p<0.001; **Supplemental Table 7**). These findings offer experimental validation for fate probabilities.

We next sought to determine whether the rate differed between inter- and intra-clonal transitions. We returned to visitation probability (VP) (**Figure 2a**) and an additional measure, conditional mean first passage time (CMFPT), to quantify the probability and rate of cells transitioning toward a single transient state. We identified the MultiLin progenitor with the highest fate to cDC, which belonged to sgRNA-5, and calculated the VP and CMFPT for every HSPC to this cell (**Figure 5g**). Cells harboring the same guide (sgRNA-5) had higher probability toward this state, with sgRNA-10 showing a similar distribution. Meanwhile the sgRNA-4 and sgRNA-9 possessed marginally lower probability. Meanwhile, HSPC populations have relatively quicker transitions to MultiLin but with a lower visitation probability. A MANOVA was conducted on the guides for both cell types and statistical significance was found (Pillai value 0.55, p-value =2.2×10^−16^; **Supplemental Table 8**). We found the MultiLin sgRNA-5 cells were significantly different from all groups (p<0.001). Finally, we sought to determine if clones had preferential visitation to specific cell population, and devised a metric termed weighted destination time (WDT). We ranked WDT for the erythroid and lymphoid lineages, and identified a diametric opposition between the lineages across the top 10 clones (*ρ* = −0.745, p-value = 0.018) (Figure 5h). Collectively, these results suggests that cell-cell transitions are more likely to occur within a clonal lineage at both the terminal differentiation state, and transitory states in between. We anticipate that these computational approaches will provide quantitative metrics to complement advances in hematopoietic stem cell [25] and leukemic cultures [26, 27] that are more broadly capable of representing stem cell biology and properties of self-renewal.

## DISCUSSION

Applying jump diffusion on the cell-cell network allows for an expanded number of neighbors of for each node over diffusion methods. This is due to the adaptive probabilistic neighbor’s search introduced. Allowing more neighbors increases the odds of capturing discontinuous processes. Conceptually, the closest related work to introducing jumps within a network is through a strategy called “lazy-teleporting”[28]. Instead of finding new edges within the network to act as jumps, we use jumps to tune a transition probability matrix within PC space. The jumps allow us to identify discontinuous processes in PC space for given cellular states. These processes can determine relevant gene programs. We demonstrate identifying these programs by splitting the positive and negative genes along a jump identified PC and perform gene Ontology. Future work will be aimed at more robust identification of gene signatures for specific processes, and tying explicit jumps with transcription factor activity. An example of a method to incorporate in the future is developing distributions of jump-related genes through gene trajectories to identify new genesets associated with eigen cells[29].

Most trajectory analysis approaches aggregate all cells together across genotypes within a unified representation, permitting transitions into states that are not biologically accessible within a given system, such as populations that are absent or experimentally ablated. When dealing with absorbing Markov chains, it is imperative the absorbing states are obtainable for the respective genotypes. We propose to handle this by decoupling the network to their respective replicates, and performing batch correction on their fate calculations. Batch correction has been developed for pseudotime, but not for fates[30–32]. These batch corrections account for different effect size and cell samplings of replicates. Corrected fates more faithfully represent their biological replicates while ensuring absorbing states are contained within each replicate. Our modular framework uses fate calculations to identify preferentially biased cells for particular lineages. This is performed by using lineages and self-membership to determine an outlier score compared to the pseudobulk for a particular cell population. This method foregoes the need for arbitrary cutoffs on fates alone as it provides an adaptive policy. The biased cells allow for novel discovery towards the mechanisms of a particular fate.

Hypothesis generation tools have been designed comparing pseudotime and lineages to differential gene expression such as Tradeseq[33] and condiments[30]. We focus on generating hypothesis tools through fates, specifically in the context of preferentially biased cells. We use an empirical CDF outlier detection to identify biased cells as categorical variables. We demonstrate the use of bias cell detection through a classical development model of GMP differentiation along monocytic and neutrophil lineages. Although subclustering methods can identify cell subsets, our outlier detection approach offers the advantage of semi-supervised labeling guided by lineage fates selected by the user. A limitation of our approach is it relies on identifying cells that are heavy tailed compared to the bulk. If the bulk of cells have a high affinity/fate for a particular lineage only the outliers will be registered as biased. Furthermore, fates are aimed at identifying terminal states, and largely ignore intermediate ones of interest. We introduce visitation probability and weighted destination time to handle potentially underrepresented biased cells and evaluate their progression through intermediate progenitors. These metrics allow us to determine how quickly and likely a cell is to visit an intermediate cell state before reaching absorbing states. We demonstrate their application in the context of biased cell calls, where biased cells have a higher probability and faster travel times than nonbiased cells.

We identify the underlying mechanisms of what makes cells biased. We do this through performing differential gene expression and transcription factor activity. While we demonstrate the use of bias cell detection on known myeloid differentiation lineages, we envision the utility as a hypothesis generation tool. Our outlier detection technique allows for binning cells by nonbiased, uni-biased, and multi-biased to recover unique characteristics within a given population for lineages. This allows for understanding the current state through RNA expression, and nominating testable mechanisms through TF activity. We fit models pairing TF activity to fates which further inform which TFs are synergistic and antagonistic to particular lineages. This has potential to impact perturbation studies where nominated TFs can readily be performed on functional assays. A limitation we face is the reliance of existing gene-TF networks as our approach does not nominate new candidates for gene TF interactions. This is left to future work along with approaches to determine the efficacy of the nominated candidates. Overall, we have introduced novel mechanisms that improve exploratory analysis within scRNA.

## ACKNOWLEDGEMENTS

We are grateful to members of the Bowman Lab and Dr. Dana Silverbush for their discussion of the work. This work was supported by the Leukemia Research Foundation, American Society of Hematology, V Foundation and the National Institute of Health (UG1CA233332, R37CA226433).

## AUTHOR CONTRIBUTIONS

M.B. and R.L.B. conceived and designed the study. R.B., and R.L.B. designed and executed lentiviral experiments. R.V.V. and S.M.S. provided critical reagents for lentiviral barcoding. M.B. and V.S. performed all computational analysis. R.B., V.S., R.V.V, S.M.S, M.T., and J.J.T. provided critical discussion on experimental design and crucial datasets. R.L.B. supervised the study. M.B. and R.L.B. wrote the manuscript, all authors reviewed and commented on the final manuscript.

## METHODS

### Lentiviral production and leukemic cell infection

To produce lentivirus, jetPRIME-mediated transfection of HEK293T cells was performed using 1.8 ug of psPAX2 helper plasmid and pMD2.G plasmid for every 2.7 ug of CROP-Seq barcoding library plasmid[20]. One day after transfection, cells were washed once with PBS and media was changed. Conditioned media was collected 48 hours later, pelleted to remove cellular debris, and filtered through a 0.45um filter. Virus was concentrated 100X by ultracentrifugation, resuspended in StemSpan (StemCell) and frozen at −80C. Whole bone marrow from leukemic *Rosa26:FlpoERT2, Npm1*^*c-Frt*^*-Flt3*^*ITD-Frt*^ mice was thawed and cultured for 1 week in StemSpan with 50ng/ml of SCF and 10ng/ml IL3. Lentiviral infection was performed with 8 ug/mL polybrene and 10 mM HEPES. These cells were cultured in StemSpan media with 1% PenStrep containing 50 ng/mL murine SCF and 10 ng/mL murine IL3 in a non-TC treated 12-well plate. 24 hours after infection, cells were collected from the plate and pelleted at 300 xG for 5 minutes in a centrifuge kept at 4°C. Media was removed and fresh media with the same composition described above was added to resuspend and replate the cells. 24 hours after media change, GFP-expressing cells were sorted and isolated on a Sony SH800S Cell Sorter. 10,000 cells were collected from the sorter.

These cells were placed back into media with the same composition described above for 9 days. After 9 days, the cells were collected and prepared for Chromium GEM-X Single Cell 3’ sequencing to determine the transcriptional profile of the clones.

### Defining pseudotime kernels in semi-supervised fashion

Our workflow begins after cell cluster identification through the Seurat package[34]. We allow for different kernels to be used to represent the cell state, specifically to order them along a pseudotime. We allow supervised signatures through either custom gene sets curator by users, or through curated sets with msigdbr package[35], such as Hallmark gene sets like Hallmark Brown Myeloid Up or Hallmark Inflammation response. After genesets are determined Seurat’s built in Add Module Score function is used to score each individual’s cell as a pseudotime (which we will denote as *T*). Alternatively, if users want to rely on a semi-supervised signature to derive genesets, we utilize Seurat’s FindAllMarkers function to find the top *g* ∈ *G*) upregulated genes for specified terminal clusters. The semi-supervised signatures then are placed into the Add Module Score function to get a score for each unique cluster specified.

When multiple pseudotimes are used to reflect the same criteria such as differentiation, we apply an entropic efficiency metric to find a single pseudotime. As an example, let us assume we have a matrix *T*^*p*×*τ*^ which contains cells as the rows and their components of for pseudotime on the columns. The pseudotime measures only account for terminally differentiated cell populations. We first row normalize the scores and formulate a matrix, 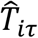.

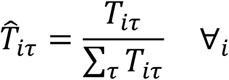

We then calculate the entropy of each cell through a standard entropy equation below:

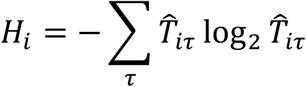

We then produce a maximum entropy test condition which is a uniform distribution for the number of possible pseudotime components, *H*_*max*_. Then we can determine the entropic efficiency, *U*_*i*_ by:

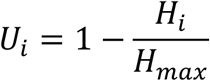

When cells hold a uniform distribution for the signatures, they represent noncommitted cells to a particular lineage. This would result in in low efficiency score which we interpret at as an early progenitor population. As the score becomes higher, cells hold a lower entropy score due to committing to a single lineage, which we interpret as a more differentiated population.

### Transforming data with cosine similarity

Cosine similarity is a measure that better encaptures clusters of high dimensional data because it is a directional measure. This is preferable to standard Euclidean distance (a magnitude based measure) as means to understand how far apart cells are to one another. Cosine similarity has been used routinely for batch correction[36]. This is particularly useful when considering a set of principle components (*d* ∈ *D*) which can carry a different significance or variance explained thus different scaling for each dimension (which we denote as *λ*_*d*_). Transforming the standard principle components, 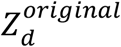 to their cosine equivalent, 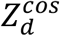is done in the following equation:

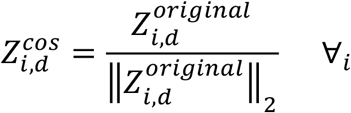

This measure improves similarity between cell types as each cell within a cluster should hold similar directionality despite differences in magnitude.

### Model fit with jump-drift-diffusion

#### Identifying eigen cell states

We rely on a synthetic eigen cells, conceptually similar to meta-cells[37], to determine our model regression for a jump-drift diffusion stochastic differential equation. We determine eigen cells by grouping all cells to their respective cell cluster, *k* ∈ *K*, then finding the average of 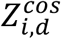.

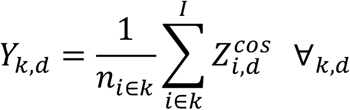

The eigen cells, *Y*_*k,d*_, allow for the model to mathematically account for directed drifting to a particular cellular state.

#### Performing forward jump-drift diffusion model fit

Traditional diffusion models follow the stochastic model for cell transitions, where *X*_*t*_ represents the state of a cell at timepoint t, *σ*_*k,d*_ represents a band of nearest neighbors is often described as a heuristic, and *d W*_*t*_ represents the Brownian motion.

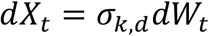

In our case, we treat *X*_*t*_ as 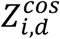and thus *dX*_*t*_ is the state transition from 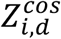to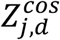. The above stochastic equation along with the definition of *dX*_*t*_ allows for diffusion probability density function to be designed as follows:

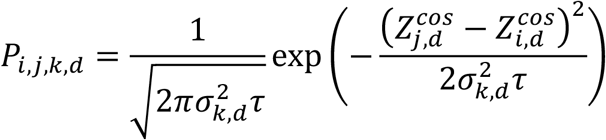

The *τ* represents the timestep (delta in pseudotime) between cell *i* and *j*. The above equation is then used to determine the affinity between cells, where they are combined to build a cell-cell network. These are commonly known as diffusion maps popularized by destiny[1]. Strategies have used temporal scRNA datasets to fit the diffusion models through neural networks[38, 39], optimal transport[6, 40], and Markov chains[41, 42]. We adapt our model for a Markov chain by introducing jumps without the power of temporal scRNA snapshots. Jumps and discontinuous processes in latent space have been described before in the context of scRNA[43].

Our model includes jump terms with an augmented stochastic differential equation[44] that is written as:

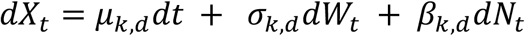

The drift term, *μ*_*x*_ represents a deliberate weighting for the current cell, 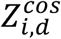, and distance to an eigen cell, *Y*_*k,d*_:

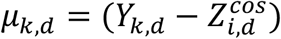

Meanwhile, *σ*_*k,d*_ is a variable to describe the diffusion of the stochastic process. β_*k,d*_ represents an additive jump process which contains three variables, 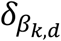representing the rate at which a jump occurs, 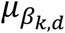 the magnitude of the jump, and 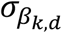 the variance on the jump magnitude. All four of these variables are to be fit through a maximum likelihood estimate. The following transition probability equation follows a modified Poisson Distribution:

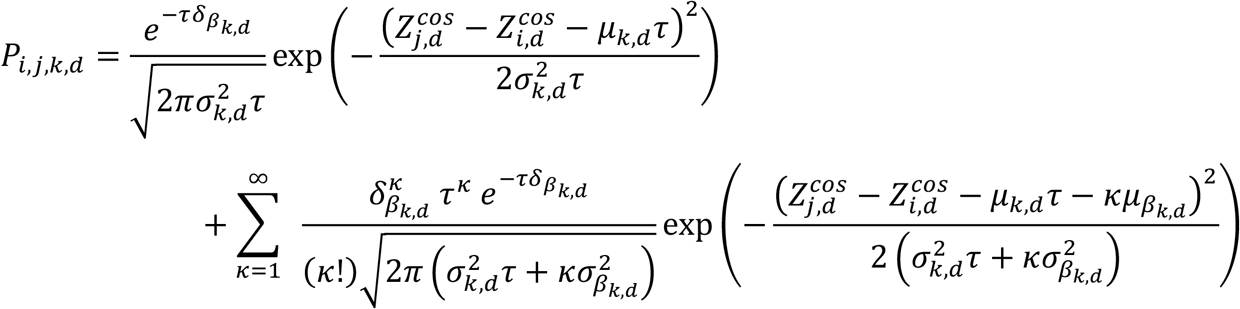

Afterwards, a series of models are fit to the data through a maximum likelihood estimate via negative log likelihood estimates as follows:

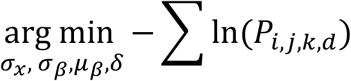

The optimization above is a constrained optimization where the data is fit using nloptr R package, where afterwards the variables are then put into the modified Poisson distribution to achieve the *P*_*i, j, k, d*_. All data is used to fit the model to determine the variables. The output probabilities for each model fit are then normalized across each principle component as follows:

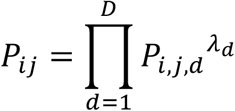

Each principle component carries a different significance of information, thus we scale the likelihood estimates together *λ*_*d*_. This allows the probability of a transition between cell M and *i. j* unique attribute of our cell-cell network compared to others including PAGA[45] is it is inherently not a guaranteed symmetric matrix, which may better reflect actual biological processes (i.e. more difficult to de-differentiate than differentiate). Heuristic cutoffs can then be applied to this matrix before establishing a Markov chain.

#### Establishing Markov chain properties

When the transition probability matrix (TPM) has been established, 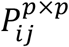 is a square matrix which contains rows *i* ∈ *p* and columns *j* ∈ *b*. We make the model follow Markovian properties by row normalizing the matrix:

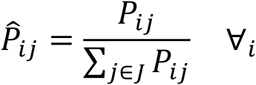

The normalization forces each row to sum to 1, thus forcing a transition action to produce an outcome. Afterwards, we can augment our Markov chain into an absorbing Markov chain by partitioning it into block format. The partition is reliant on selecting cells to act as absorbing states, in that, these are states once entered, cannot leave. Cell states that are not absorbing states are referred to as transition states. The absorbing cells will be denoted as m, while transition cells will be labeled as n. The total number of rows (and columns) is *p* = *n* + *m* which accounts for all the cells. The blocks can be defined as follows: 1) Transition probability for transition states to other transition stated with be denoted as *Q*^*n*×*n*^, 2) transition probability for transition states to absorbing states will be denoted as *R*^*n*×*m*^, 3) absorbing states cannot transition to any other state, thus are augmented with a zero matrix 0^*m*×*n*^, and 4) absorbing states can only stay in their own state, this is represented by an identity matrix *I*^*m*×*m*^. We can rearrange the rows and columns of the TPM to fill in these four blocks as follows:

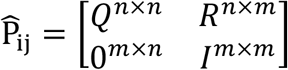

#### Finding cell fate and visitation probability as a lineage representation

The block representation allows us to determine a closed form solution to random walks by solving a fundamental matrix *N*^*n*×*n*^ by inverting a subtraction of the identity matrix.

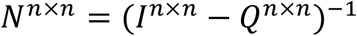

*N*^*n*×*n*^ represents the expected number of visits before being absorbed from transient cell i to transient cell j. We then use *N*^*n*×*n*^ to obtain the fate matrix, *B*^*n*×*m*^. The fate matrix is the expected number of transient visits by the probability to transition from a transient state to absorbing state, *R*^*n*×*m*^.

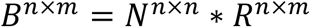

To extend the *B*^*n*×*m*^ to all cells, we append *B*^*n*×*m*^ to include an identity matrix to account for each absorbing states, *I*^*m*×*m*^, to fully realize *B*^*p*×*m*^. To determine lineage specific fates, we first use a priori information to define cells *j* into sets of lineage clusters, *l* ∈*L*. The probability of being absorbed into any cell assigned to a cluster n is independent of one another and achieves the same macroscopic view of reaching that cluster, effectively allowing us to treat them as an “or” operator which we can sum across sets.

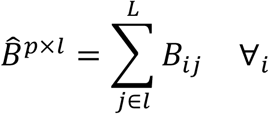

However, a drawback of this approach is the sensitivity to determining the appropriate number of absorbing states and preserving imbalanced datasets. A solution for this is to perform batch correction on the fate calculations to account for effect size. This is done by building a linear model using a standard lm function in R. We build a full model that accounts for replicates as a factor, and a reduced model that ignores the replicates. The common factors between both models are the assigned cell cluster and lineages. Subtracting the two fitted models from each other produces a corrected model. This is reminiscent of a similar technique seen in Bulk RNA sequencing known as ComBAT[9, 10].

Alternative to fates, we rely on visitation probability, *H*^*n*×*n*^, through the following equation:

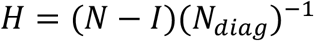

Visitation probability allows us to determine which cells are traversed before being absorbed at all. Fates inform solely about terminal state absorption. We use. in conjunction with 2 other measures to determine an alternative measure for biased cells. We rely on a measure called Conditional Mean First Passage Time (CMFPT), which is the average first visit to state*j* when starting at state *i*. To calculate this measure, Γ_*ij*_, we effectively iterate over each cell and treat it as the only available absorbing state, such that a column vector is formed by 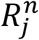, and then we reacquire matrices B and N.

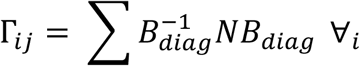

We further developed a heuristic called *weighted destination time, D*, to inform whether cells of interest are uniquely adept at visiting certain populations over others. First, based on 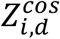, we soft cluster each cell for their respective cell type cluster through Expectation Maximization to obtain membership probabilities, *M*^*nxk*^, through the posterior probabilities. A strategy inspired by transition scores of MuTrans[46]. A diagonal matrix represents the size of each cluster,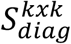, is also constructed to account for population imbalances:

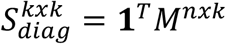

Afterwards, we perform batch correction on both the *H* and Γ following the same strategy as the fates by swapping the lineage factor for the cell j’s cluster. Then element wise division of the visitation probability and the CMFPT is performed and row normalization to represents the relative flux each of cell, *Â*_*ij*_.

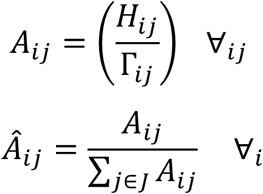

Obtaining the weighted destination time combines the cell to cell flux and the membership of a cell as follows:

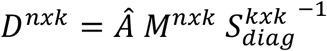

A unique aspect of weighted destination time is this measure allows for condition specific flux to clusters. Let lineages be a one hot encoder for each cell, *L*^*nxl*^, and a diagonal size matrix,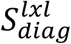, to account for population imbalance is created by through the same mechanism as 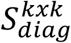. The lineage specific weighted destination time is then completed as:

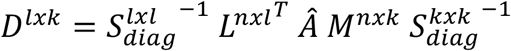

While this allows for a size correction of lineages and clusters, we aim to handle general connectivity differences within the network to evaluate lineage skewing. A global connectivity can be considered, and an expected weighted destination time can be made by treating all cells as a single lineage. Afterwards a log2 fold change can be taken between *D* ^*lxk*^ and the expected weighted destination time to identify enrichment of specific lineages.

### Identifying preferential lineage biased cells

First, we group cells by their cell type classification to determine preferential biases towards a cluster. We then use cells from this cluster to determine preferential biases for each lineage independently. To find preferentially biased cells we rely only on lineage values from aggregated fate. We build an empirical cumulative density function for each lineage of the users’choice, 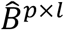.

A right tailed cumulative density function, 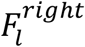, is built to find the values skewed toward a higher probability to be classified as preferentially biased for each individual lineage. Outliers scores for each cell in the lineage, *O*_*i,l*_, are determined as the natural log of the cumulative density function.

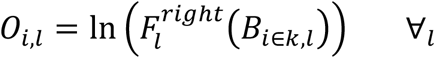

We then determine the skewness of the distribution to ensure an appropriate scaling of the score can be attributed as below:

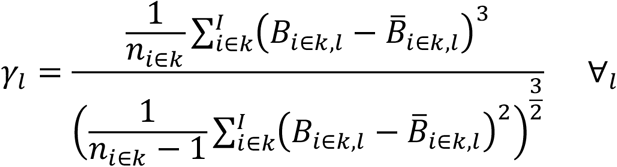

We then calculate the skewness of the membership following the same formula.

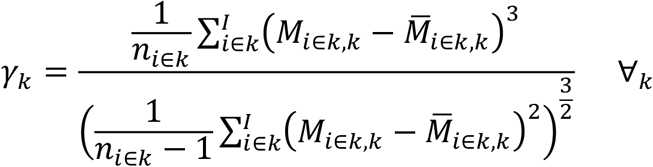

Next, we aggregate a cell’s total outlier score by summing across their individual outlier scores with skew measures and is designed to find cells that exceed high levels of outlier score:

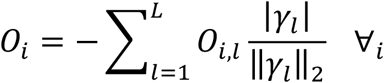

With the aggregated outlier scores, we augment them to through a robust quantile regression filter. This is conducted by using a Huber loss function with the acceptable bounds determined by the median absolute deviation of *O*_*i*_. The entirety of the piecewise Huber loss function used is shown below:

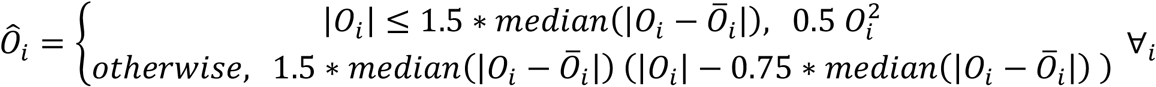

Once the robust regression is performed, a cutoff value to determine inlier vs outlier cells is through a standard Tukey’s rule is applied:

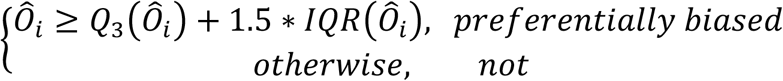

These binary classifications are then used to determine preferentially biased cells in downstream analysis.

### Transcription factor network and inferencing

We rely on the decoupleR package for inferring transcription factor enrichment[13]. This code base uses a transcription factor network model developed by swissregulon for interaction weights between transcription factors and genes. The interaction weights are denoted as *ω*_*gt*_, with *g* representing the set of genes and *t* representing the set of transcription factors. The RNA expression matrix for each cell is represented by *G*_*ig*_ where *i* is each cell, and g is each gene. Each transcription factor is fit with a univariate linear model to predict observed gene expression based on this interaction weights:

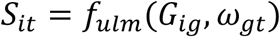

After the model is fit, a student t distribution is used to score transcription factor activity (*S*_*it*_). Positive values indicate active transcription factor and a negative value inactive.

We use the transcription factor information along with the preferentially biased calls from *ô*_*i*_ to find differential activity within cells per cell type cluster. Further, we also have a built-in function that allows the comparison of different conditions (i.e. age or order of mutation) for the transcription factor activity as well.

### Tying transcription factor activity and lineages together

Once transcription factor activity is inferred for each cell, *S*_*it*_, a multivariate linear model is fit with lineage probabilities, *B*_*ik*_, and transcription factors using a standard *lm* function in R.

Transcription factors are the independent variables while the lineages are the response variables.

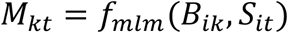

The output of the model produces scores and p-values that signal which transcription factors are active and inactive with significance for each lineage, which are then used for hypothesis generation. We have built in functions that compare the transcription factors to different lineages in a pairwise manner to determine whether transcripts are synergistic or antagonistic. These directly are used for hypothesis generation to determine candidate transcripts for functional assays.

### Identifying jump programs within principle components

#### Tying the jump programs to cells of interest

Jumps help nominate potential cell type groups that contain discontinuous processes, more specifically, they aid in nominating which gene features within a principle components should be investigated. We can first identify discontinuous processes for jumps via the model fits for each PC. A jump, essentially signifies a significant deviation in which diffusion can not obtain alone, is necessary to reach an eigen cell at a given pseudotime. As an example, let us turn our attention to the GMP jump in PC4 as seen in **Figure 3b**. Since our model is built on the propagating the earliest pseudotime state, in this instance *U*_*i*_= 0, a jump of −0.45 would be required to reach the GMP eigen state within the allotted pseudotime.

A higher absolute magnitude means a significant deviation from diffusion must occur for the initiating cell to propagate to an eigen cell. Our model essentially fits data by considering an initiating state to all other states toward a particular eigen cell at a particular pseudotime. A higher absolute jump value indicates a more significant action is needed from the initiating state (earliest pseudotime value, in this instance *U*_*i*_ = 0), to achieve the eigen cell state. An example of jumps vs no jumps is shown occurring in the model fit is shown in **Figure 3b**.

#### Extracting relevant gene ontology pathways from jumps with JumpROPE (relevant ontology pathway extraction)

While some strategies aim to correlate gene patterns through generative models[47], we rely on known gene signatures to explain jump programs. An additional aspect of interest is identifying genes associated with jumps given cell populations and principle components. After the jump models are fit for each individual cell population along principle components, we can identify whether a jump is used to fit the data as shown in **Figure 3c**.

Afterwards, we can identify each PC that has an associated jump, and extract the strongest contributing genes to those PC_s_, *ϕ*_*gd*_. Specifically, we do this by first centering, scaling, and normalizing each principle component feature embedding, *Z*_*gd*_, where a jump is identified:

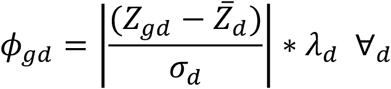

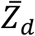 is the mean value of the principle component *d*, and *σ*_*d*_ is the standard deviation. We are not interested in how these genes are impacting the principle component, but rather their rank which is why the absolute value is used. Further, each principle component is scaled by their importance *λ*_*d*_. Then we find the genes that are outliers through the standard Tukey equation where *Q*_3_ represents the third quantile:

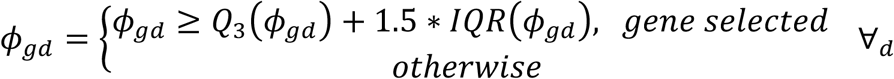

Afterwards we allow a cutoff of the top n genes, which we further split into their positive and negative loadings by loading them back into the initial gene loading embedding. Once the loadings are split, GeneOntology analysis is conducted through the TopGO package[48]. An example of the top 15 genesets is shown in **Figure 3c**.

## CODE AND DATA AVAILABILITY STATEMENT

The opensource package is available for download at https://github.com/namwob44/SupeRJump/.

A link for the specific codes and data used in this study will be made available upon request only prior to publication.

